# Conditioned overconsumption is dependent on reinforcer type in lean, but not obese, mice

**DOI:** 10.1101/2023.12.31.573797

**Authors:** Darielle Lewis-Sanders, Sebastien Bullich, Maria Jose Olvera, John Vo, Yang-Sun Hwang, Sarah A. Stern

## Abstract

Associative learning can drive many different types of behaviors, including food consumption. Previous studies have shown that cues paired with food delivery while mice are hungry will lead increased consumption in the presence of those cues at later times. We previously showed that overconsumption can be driven in male mice by contextual cues, using chow pellets. Here we extended our findings by examining other parameters that may influence the outcome of context-conditioned overconsumption training. We found that the task worked equally well in males and females, and that palatable substances such as high-fat diet and Ensure chocolate milkshake supported learning and induced overconsumption. Surprisingly, mice did not overconsume when sucrose was used as the reinforcer during training, suggesting that nutritional content is a critical factor. Interestingly, we also observed that diet-induced obese mice did not learn the task. Overall, we find that context-conditioned overconsumption can be studied in lean males and female mice, and with multiple reinforcer types.

## 1. Introduction

Feeding is defined as a complex motivated behavior, and although the primary driver of food consumption is the homeostatic maintenance of energy balance, many other factors can influence when, what and how much food is consumed (Azevedo et al., 2022; Berthoud et al., 2017). Associative learning is one such factor (Thibault, 2010).

Experimental evidence that associative learning can drive food consumption was originally demonstrated by Henry Weingarten in the 1980’s (Weingarten, 1983), where he showed that hungry rats who were given access to food paired with a specific tone had a lower latency to consume following the tone, even once rats were sated. Furthermore, rats consumed over 20% of their total daily caloric intake in response to this conditioned cue.

Petrovich and colleagues extended these findings in a number of studies (Cole et al., 2015; Holland et al., 2002; Petrovich, 2011; Petrovich et al., 2007, 2012; Petrovich & Gallagher, 2007), demonstrating a role for amygdala, hypothalamic and prefrontal brain regions in regulating overconsumption behavior in rats. The majority of these studies used a discrete auditory or visual cue to pair with food delivery during the training, although one study demonstrated that contextual cues were also effective at eliciting overconsumption (Petrovich et al., 2007). One other study adapted the task to mice (Walker et al., 2012)

Recently we developed a modified version of this task for mice in which contextual cues are utilized to elicit the conditioned overconsumption response (Stern et al., 2020). In this task, mice are food deprived overnight and then given access to chow in a novel context. After a number of training sessions, mice are returned to ad libitum feeding for 2-4 days. They are then placed back into the context and food intake under satiation is measured. Controls can be conducted either between-subjects (Stern et al., 2021) or within-subjects (Stern et al., 2020). In the between-subjects condition, a second group of control mice is habituated to the novel context and fasted in their home cage at paired times with the experimental group, ensuring that they are exposed to the novel context and also experience equivalent levels of fasting. They are subsequently also offered food in the novel context during testing. Because they do not receive food paired with the context during training we refer to this group as “CTX-,“ and the group trained in the novel context as “CTX+.” When tested within-subjects, each mouse is habituated to two novel contexts (CTX+ and CTX-), and then only the CTX+ is used for training sessions. During testing, mice are re-exposed to both contexts, counterbalanced, with food available. In both cases, this task results in increased consumption in the CTX+ compared to the control mice or to mice tested in the CTX-.

Our initial investigation tested the effectiveness of the task using massed or spaced training, as well as the role of positive and negative reinforcement as drivers of the response (Stern et al., 2020), but it left open a number of experimental questions. First, does the task work equally well in male and female mice? Our original study tested only male mice, but testing in female mice is important for future studies. Second, we originally tested mice using cocoa-flavored chow pellets as the unconditioned stimulus, but it has previously been shown that the reinforcer type used as the unconditioned stimulus in appetitive Pavlovian conditioning can have an effect on learning (Stanhope, 1989). Therefore, we ask whether non-chow foods support learning of the overconsumption task? Third, are animals with experimentally-induced obesity predisposed to this type of conditioned overconsumption? Here we address these questions with a series of experiments, and find that although the task works equally well in male and female mice, not all food reinforcers are able to support learning, possibly due to their palatability or nutritional content. Moreover, we find that experimentally obese mice are unable to learn the task, as supported by previous literature in learning and memory tasks.

## 2. Methods

### 2.1 Mice

The experiments in this study were approved by the Max Planck Florida Institute for Neuroscience Animal Care and Use Committee (Protocol #20-007) and were in accordance with the National Institute of Health Guide for the Care and Use of Laboratory Animals. Male and female C57BL/6J mice (Jackson Laboratory, 000664) were 8-20 weeks at the start of behavioral testing. Mice were group housed on a 12-hour (h) light/dark cycle with *ad libitum* access to water and standard mouse chow (PicoLab Rodent Diet), except when fasted as noted below. Diet-Induced Obese (DIO) animals were generated by giving C57BL/6J mice *ad libitum* 60% high-fat diet (Research Diets, D12492) starting at 6 weeks old. Mice were fed on high-fat diet (HFD) for at least 12 weeks before being used for behavioral procedures. Mice were individually handled 1-2 minutes for 3 days prior to the start of any behavioral procedures.

#### 2.2 Context-conditioned overconsumption

Context-conditioned overconsumption was performed as described previously (Stern et al., 2021), with modifications described below.

##### Behavioral acclimation

In order to prevent neophobia, animals were given some of the test-food that they would be exposed to during the experiment after each handling session. Ten chocolate-flavored Precision Pellets (Bio-Serv, F05301), sucrose pellet (Bioserv, F07595) or chocolate sucrose pellets (Bioserv, F0025) were placed on the homecage floor on day 1 of handling and 5 pellets were placed on the homecage floor on subsequent days. Animals given 60% high-fat diet (Research Diets, D12492) in the task were given pellets in the same procedure as above. Animals that received 50% Ensure (Abbot Laboratories, Chicago, Ill) or 10% sucrose (Fisher Chemical, S5-500) in the task were given 3mL in their homecage on each day of handling.

##### Room acclimatation

Animal cages were placed in the experimental room 30 minutes prior to the beginning of the task phase. All behavioral experiments were performed in the morning starting around 9:30am.

##### Habituation

Animals were placed in a novel context that has a distinct shape, color, bedding, and location from the homecage. Animals were placed in the chamber for 20 minutes and then returned to their homecage. At the end of context habituation, cages were randomly separated into two groups, CTX+ and CTX-.

##### Training

All mice were fasted 18hrs prior to training and the home cage was changed to prevent animals from eating food dust. After fasting, animals in the CTX+ group were placed in the conditioning chamber with 2.0 grams of pellets for 30 minutes. Animals being trained with HFD were given 3-4 grams in the conditioning chamber. Animals given 50% Ensure in the task were given 10 mL. As a control, the CTX-animals were kept in their homecage during the CTX+ group training period. Following training, pellets were weighed, and mice were returned to their homecage. One hour after the end of the training session, the mice are given food back *ad libitum.* In the pre-exposure HFD group, animals in the CTX-group were also given HFD for 30 minutes in their home cage one hour after being provided with food in the home cage. This protocol is repeated for training session 2 and 3 separated by 48h.

##### Testing

After the last training day, animals were fed *ad libitum* for 48h prior to testing. Both CTX+ and CTX-animals were placed in the chamber for 20 minutes with measured diet, as indicated above. After testing, the diet was weighed, and the mice were returned to their home cage.

#### 2.3 Conditioned taste aversion (CTA)

CTA was performed as described previously (Olvera & Miranda, 2019), with modifications described below.

##### Habituation

Mice were habituated to the licking box chamber for 30 minutes and were then water deprived for 12h. To establish basal liquid consumption, they were habituated for 3 days to have water access for 20 min/day (around 12:00 and 14:00 h) from a graduated bottle in the licking box chamber.

##### CTA Training

Sixteen hours before the CTA training, mice underwent water deprivation. During the 20-minute training session, mice were presented with a new flavor: either a 10% sugar solution (Fisher Chemical, S5-500) or 50% chocolate Ensure (Abbott). Thirty minutes later, mice were intraperitoneally injected with 0.30 M LiCl (10 ml/kg), a substance known to induce malaise. Water access was reinstated after the intraperitoneal injection. Post-LiCl injection, mice were monitored in their homecage, and at 7 pm, water deprivation was initiated.

##### CTA Testing

The test was conducted 24 hours after the training. Sixteen hours prior to testing mice were water deprived. During testing all mice were once again exposed to either the sucrose or Ensure solution for 20 minutes. The reduction in sugar or Ensure consumption during the retrieval test was compared with the consumption during the acquisition phase. Consequently, the strength of CTA was calculated as the percentage of sucrose or Ensure intake during testing relative to CTA training.

#### 2.4 Immediate early gene mapping

The context-conditioned overconsumption protocol was conducted as described above with the following changes. Thirty minutes after the test phase, both groups of mice were perfused with 4% paraformaldehyde (PFA). Brains remained overnight in 4% PFA before being transferred to phosphate-buffered saline (PBS). Mice brains were serially sectioned at 50 µm into three individual wells Sections were washed three times for 15 minutes each, using PBST (PBS-0.2% TritonX) then blocked with 3% goat serum in 3% BSA with 0.05% sodium azide. Sections were then incubated 1:500 in primary cFos antibody (Cell Signaling, Danvers, MA #2250) for 48h at 4°C. Sections were then washed three times in PBST, then incubated 1:1000 in goat anti rabbit AlexaFluor 488 secondary antibody (Thermo Fisher Scientific, Waltham, MA, #A11008) at RT for an hour. Sections were washed again in PBST 3 times and were mounted using Fluoromount-G with DAPI (Thermo Fisher Scientific, Waltham, MA). They were imaged on a BZ-X800 Analyzer Keyence fluorescence microscope (Keyence, Osaka, Japan). Equivalent sections from the different groups were imaged using the same settings. cFos density was quantified using ImageJ (Schneider et al., 2012). A threshold was applied and cells are analyzed by an experimenter blinded from which group each image belonged to.

## 3. Results

### 3.1 Context-conditioned overconsumption is inducible in both male and female mice

In this study we aimed to extend the context-conditioned overconsumption task to different reinforcers and diet in order to better understand this phenotype in sated mice. First, we compared performance in the task using both female and male C57Bl6/J mice, since the task was previously demonstrated only in males (Fig1A-E). Here we used the between-subjects version of the task (Fig.1A). The animals were first habituated to a novel context and then were divided in 2 groups, CTX-and CTX+. The control CTX-group was food deprived in their homecage for 18h, but not trained for the context-conditioned overconsumption task. The CTX+ group was trained 3 times by being given food in the conditioning chamber following an 18h fast. During training, male and female mice ate comparable amounts, with intake escalating on each day (Fig. 1B-C). (Fig. 1B; Females: Repeated-measure one-way ANOVA: Session: F(1.749,12.24)=12.94, **p= 0.0012, followed by Tukey’s multiple comparison test: *p=0.0394 : Train 1 vs. Train 2; **p=0.0079 : Train 1 vs. Train 3) (Fig. 1C; Males : Repeated-measure one-way ANOVA: Session : F(1.446,10.12)=5.835, *p= 0.0273, followed by Tukey’s multiple comparison test: **p=0.0096 : Train 1 vs. Train 2). Results are presented in kilocalories (kcal) as well as (g) for later comparison with liquid substances that are measured in millileters (mL). Following the return to *ad libitum* feeding, during testing, both groups are sated and are placed in the conditioning chamber with food available. We found that both male and female CTX+ animals eat more than CTX-animals, consuming an average of 1-1.5g of cocoa-flavored chow pellets in a 20 minute test session (Fig. 1D-E) (Fig 1D; Females: *t* test: *t*_(14)_=2.796, *p=0.0143) (Fig. 1E; Males : *t* test: *t*_(14)_=2.559, *p=0.0227). Since there was no difference between males and females in the effectiveness of the task, we decided to pursue our subsequent investigation combining data from both sexes within the groups.

**Fig 1.**
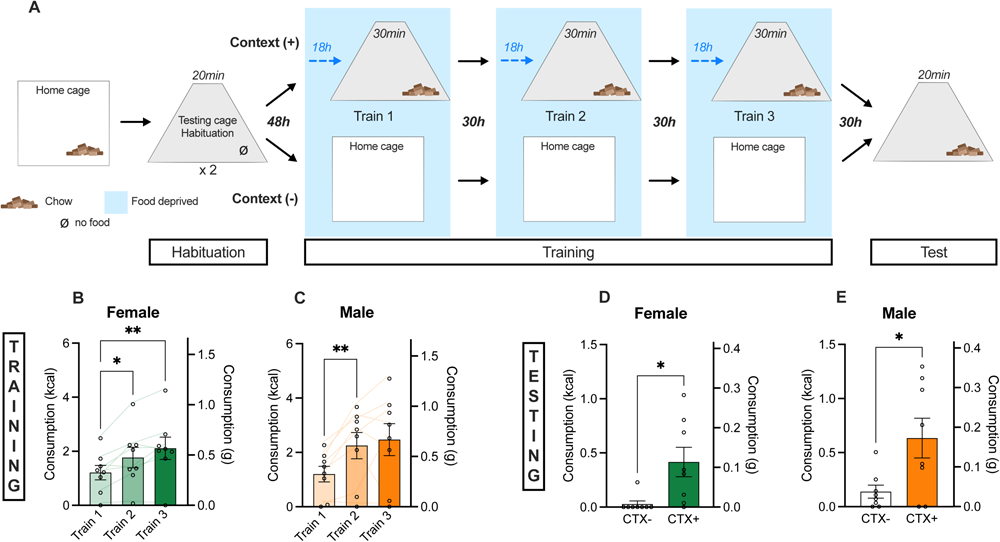
Context-conditioned overconsumption paradigm induces overeating in both males and females. (A) Experimental design of context-conditioned overconsumption paradigm. (B) Food consumption during the 3 training sessions (kcal or grams) for females and (C) males. (D) Food consumption during testing (kcal or grams) for females and (E) males. n=8/groups. t-test : *p<0.05 and **p<0.01.

### 3.2 Effectiveness of conditioned overconsumption depends on reinforcer type

We next tested whether different food reinforcers, commonly used in the field of learning and reward, can promote and/or influence context-induced feeding (Fig 2A-2I). To do so, we first tested sucrose and chocolate sucrose pellets, as well as 10% liquid sucrose. For each experiment, we habituated animals to the food after each handling sessions for 3 consecutive days to avoid neophobia during training and testing (Fig 2A). During training, all reinforcers were eaten to some extent over the different training sessions (Fig. 2B-D). However, a comparison to chow pellet consumption indicates that mice do not consume as many total calories over the training sessions when given sucrose pellets, chocolate sucrose pellets, or liquid sucrose compared to chow pellets (Supplementary Figure 1A; One-way ANOVA: F(5,58)=41.69, ****p<0.0001, followed by Tukey’s multiple comparison test: **p=0.0075 : Chow vs. Sucrose; **p=0.0058 : Chow vs. ChocSuc; ****p<0.0001 : Chow vs. LiqSu). Only the training with liquid sucrose showed a significant increase of consumption over the sessions (Fig. 2D; Liquid sucrose: Repeated-measure one-way ANOVA: Session: F(1.851,12.96)=5.408, *p=0.0213, followed by Tukey’s multiple comparison test: *p=0.0485 : Train 1 vs. Train 3) During the testing period, only sucrose pellets induced significant overconsumption in the CTX+ animals (Fig. 2F; Sucrose pellet: *t* test: *t*_(14)_=2.685, *p=0.0178), although it is important to note that the amount of sucrose pellet consumed is significantly lower chow pellets, leading us to conclude that sucrose is not a good choice for inducing context-conditioned overconsumption (Supplementary Figure 1B, One-way ANOVA: F(5,58)=6.034, ***p=0.0001, followed by Tukey’s multiple comparison test: *p=0.0365 : Chow vs. Sucrose).

**Fig 2.**
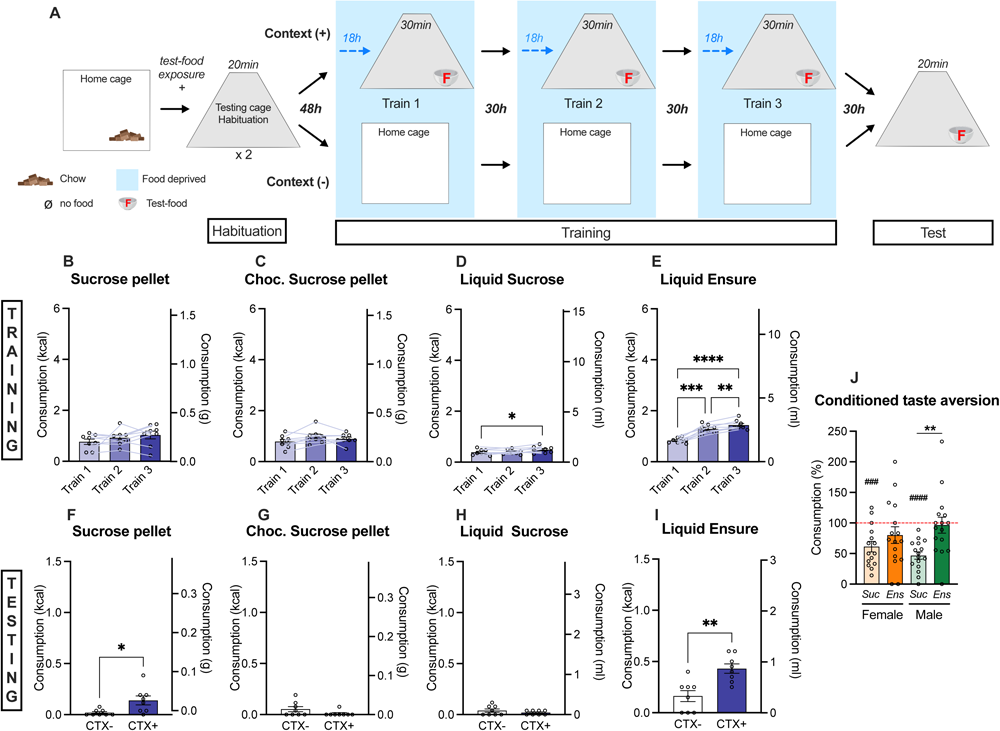
The nature of the food used as the reinforcer is important for the success of the context-conditioned overconsumption task. (A) Experimental design of context-conditioned overconsumption paradigm. The food used for the task were introduced before training session just after handling sessions to avoid neophobia. (B) Food consumption during training, while trained with sucrose or (C) chocolate sucrose pellet, (D) liquid sucrose (10%) or (E) Ensure (50%) expressed in kcal or grams. (F) Food consumption during testing while trained with sucrose or (G) chocolate sucrose pellet, (H) liquid sucrose (10%) or (I) Ensure (50%) expressed in kcal or grams. (J) Conditioned taste aversion experiment showing the percentage (%) of 10% liquid sucrose consumed during testing compared to training (testing consumption/(training consumption*100). (B-E) n=8/groups RM-ANOVA, (F-J) N=8/groups t test, (J) n=16 for Chow and HFD One-way ANOVA and (H) n=16 for M Sucrose, M Ensure and F Ensure and 15 for F Sucrose Two-way ANOVA and One sample t-test using 100% as hypothetical value. *p<0.05, **p<0.01, ***p<0.001, ****p<0.0001 and ###p<0.001, ####p<0.0001 compared to 100%.

We then tested whether liquid Ensure (50%) would be more effective at inducing overconsumption (Fig 2E, 2I). Similar to chow (Fig. 1), mice escalated intake of Ensure throughout the training sessions, and displayed significant overconsumption in the CTX+ (Fig. 2E; Repeated-measure one-way ANOVA: Session: F(1.619,11.33)=95.98, ****p< 0.0001, followed by Tukey’s multiple comparison test: ***p=0.0001 : Train 1 vs. Train 2; ****p<0.0001 : Train 1 vs. Train 3; **p=0.0062 : Train 2 vs. Train 3). During testing, CTX+ mice consumed significantly more Ensure than CTX-mice, comparable to levels seen when tested with chow pellets (Fig. 2I; Supplementary Figure 1, Ensure : *t* test: *t*_(14)_=3.811, **p=0.0019); indicating that Ensure if effective as a reinforcer in the context-conditioned overconsumption task. We therefore considered that the palatability and nutritional content of Ensure is higher than that of sucrose, conferring an advantage for this type of context-food association.

To test this hypothesis, we performed conditioned taste aversion (CTA) in which animals are trained to reduce their consumption after association with gastric malaise (Supplementary Figure 2A). We reasoned that because a negative association is learned in this task, which reduces subsequent food intake, highly palatable substances might be less likely to induce the aversive response. In other words, we wanted to test whether liquid sucrose and Ensure would induce a different response because of their palatability/nutritional content, explaining our results in the context-conditioned overconsumption task. Indeed, even though the same volume of liquid sucrose or Ensure was ingested during the training session (Supplementary Fig. 2B-C), the aversion, represented as a decrease of solution taken during the test compared to the training session, is only induced with sucrose (Fig. 2J; One sample *t* test compared to hypothetical value 100%: Female, *t*_(14)_=4.453, ###p=0.0005; Male, *t*_(15)_=8.710, ####p<0.0001). In contrast, there was no significant reduction of the consumption on Ensure during the CTA test (Fig. 2J; One sample *t* test compared to hypothetical value 100%: Female, *t*_(15)_=1.446, p=0.1688; Male, *t*_(15)_=0.2709, p=0.7902), indicating that Ensure might indeed be more palatable than sucrose. These results then led us to ask whether high-fat diet, another appetitive but also high caloric diet, would also be suitable for the overconsumption task.

### 3.3 High fat diet is effective at inducing context-conditioned overconsumption with adequate habituation

We next tested 60% high-fat diet (HFD) as a reinforcer (Fig. 3A) and, as expected, the animals escalated their intake along the different training session with a significant increase between sessions (Fig. 3B; Repeated-measure one-way ANOVA: Session: F(1.917,28.76)=15.74, ****p<0.0001, followed by Tukey’s multiple comparison test: **p=0.0025: Train 1 vs. Train 2; ***p=0.0002: Train 1 vs. Train 3). In absolute amounts, mice ate more HFD during training than chow or Ensure, demonstrating its palatability (Supplementary Figure 1A; One-way ANOVA: F(5,58)=41.69, ****p<0.0001, followed by Tukey’s multiple comparison test: ****p<0.0001 : Chow vs. HFD; ****p<0.0001 : Ensure vs. HFD). Surprisingly, however, during testing there was no significant difference of HFD consumption between groups (Fig 3C; *t* test: *t*_(30)_=1.854, p=0.0736). We also noticed that this lack of effect was driven by the CTX-animals, which ate an unusually high amount of food during the test compared to the other reinforcers that we tested previously.

**Fig 3.**
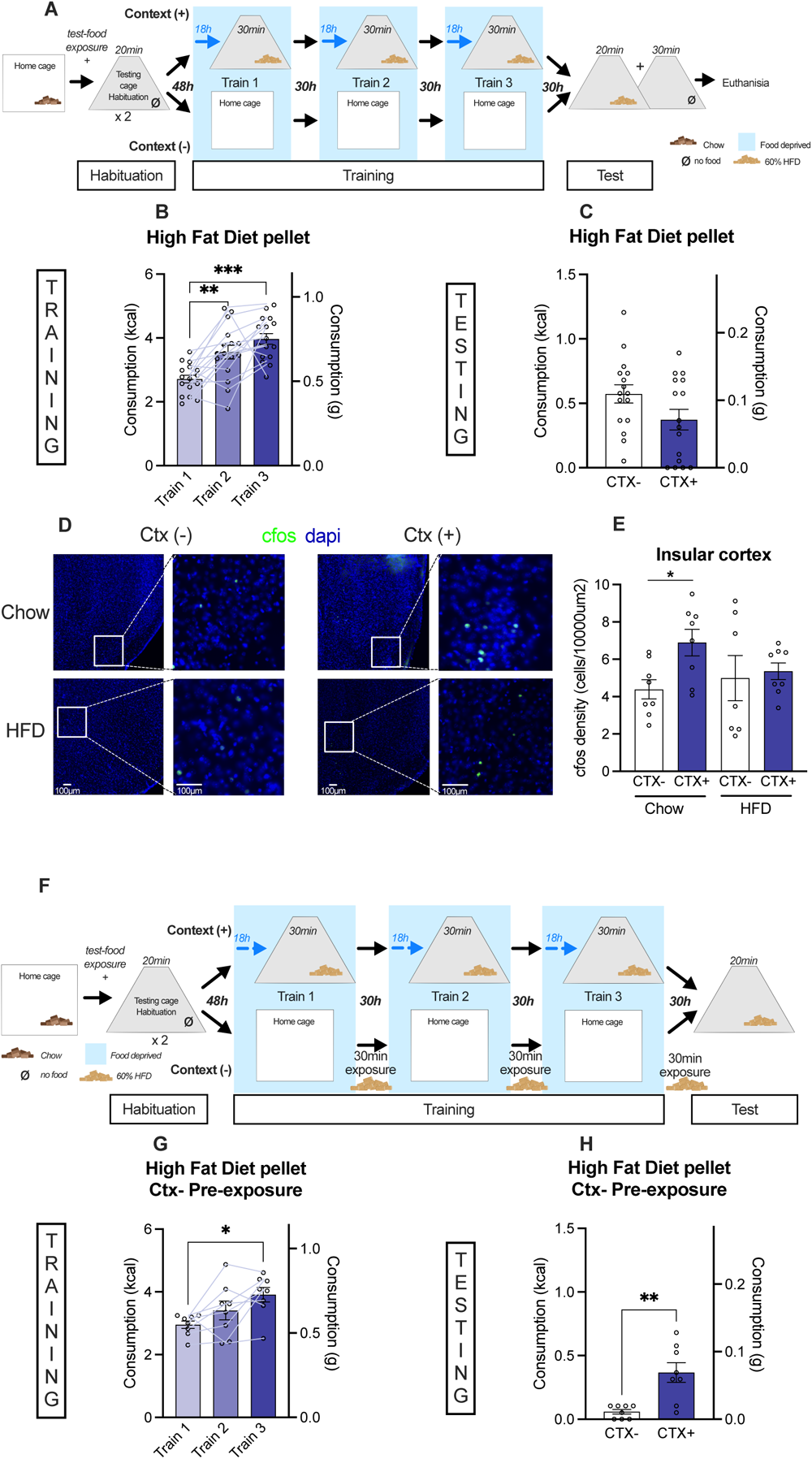
Context-conditioned overconsumption induced using high-fat diet requires adequate pre-exposure to correctly model overconsumption. (A) Experimental design of context-conditioned overconsumption paradigm. After testing, animals stayed 30min more in the context in order to euthanize the animal approximately 1h after starting testing for cFos imaging. (B) Food consumption during training while trained with HFD expressed in kcal or grams. (C) Food consumption during testing while trained with HFD expressed in kcal or grams. (D) Representative images of cFos and DAPI expression within insular cortex brain slices of Chow and HFD untrained and trained animals after context-induced overconsumption paradigm. Each group is represented by a duo of picture with an overview of the insular cortex (left) and a zoomed-in one corresponding to the square area (right). (E) Quantification of cFos expression normalized by insular cortex area expressed in cells/10000□m2. (F) Experimental design of context-IF paradigm. CTX-animals were pre-exposed to HFD (home cage) the same amount of time as CTX+ animals but not during deprived period. (G) Food consumption during training expressed in kcal or grams. (H) Food consumption during testing expressed in kcal or grams. (B) n=16/group RM-ANOVA. N=16/group t-test. (E) n=8 for CTX-Chow, CTX+ Chow and CTX+ HFD and n=7 for CTX-HFD Two-way ANOVA. (G) n=8/group RM-ANOVA. (H) n=8/group t-test. *p<0.05, **p<0.01 and ***p<0.001.

Since it has been shown that the insular cortex is activated by context-conditioned overconsumption (Stern et al., 2020) and is necessary for the association formation, we then looked at cFos expression within this structure in HFD-trained animals (Fig. 3D-E). Replicating the results from Stern et al., we found increased cFos expression within the insular cortex of CTX+ animals trained with chow. In contrast, there was no significant increase for the CTX+ animals trained with HFD (Fig. 3E; Two-way ANOVA: CTX+ vs. CTX-: F(1,27) = 3.708, p=0.0648, Chow vs. HFD: F(1,27) = 0.3906, p =0.5372, Interaction: F(1,27) = 2.048, p=0.1639, followed by uncorrected Fisher’s LSD comparison: CTX-vs. CTX+, Chow *p=0.0228, HFD p=0.7337). However, upon separating the data by sex, we found that high-fat diet impacted cFos expression in males and females differently. Indeed, in the CTX-female group, those given HFD displayed an increased cFos expression compared to their respective Chow control, while CTX-males given HFD showed a decrease compared to their respective Chow control (Supplementary Figure 4A-B).

We reasoned that the increase in consumption of CTX+ mice was either due to increased motivation of the control group or a lack of association for the trained group, so we next exposed the CTX-group to a paired amount of HFD in their homecage during the training period in addition to ad libitum access to chow diet (Fig. 3F). Interestingly, when CTX-animals had these additional exposures to HFD, they ate significantly less than CTX+ animals during testing (Fig 3H; *t* test: *t*_(14)_=3.843, **p=0.0018). Overall, these result shows that HFD is sufficient to reinforce conditioned overconsumption, but that accurate habituation to palatable/caloric reinforcers is required.

### 3.4 Diet-induced obese mice are unable to learn context-conditioned overconsumption

Because obesity may be driven by excess calories, we wanted to know if obesity may predispose an individual to conditioned overconsumption. We therefore extended the task to the diet-induced obesity (DIO) model in order to study the impact of obesity on the context-conditioned overconsumption paradigm. To do so, we fed C57Bl/6J mice with HFD for 12 weeks and we then performed the overconsumption task using the effective reinforcers described above, chow, Ensure and HFD (Fig. 4A). Surprisingly, DIO animals did not eat chow (Fig. 4B) or HFD (Fig. 4D) during training sessions, though they did ingest limited amounts of Ensure with an increase over the sessions (Fig. 4C; Repeated-measure one-way ANOVA: Session: F(1.479,10.36)=10.52, **p= 0.005, followed by Tukey’s multiple comparison test: **p=0.0088 : Train 1 vs. Train 3). As expected, during testing, there was no overconsumption phenotype for the CTX+ animals when trained with chow (Fig. 4E) or HFD (Fig. 4G), since they did not eat during training. Despite their consumption during training, there was also no significant difference between groups when trained with Ensure (Fig. 4F). The task was also not effective with sucrose, as in lean mice (Supplementary Figure 3A-B). Overall, these experiments show that DIO animals were not able to learn the overconsumption task and may require different experimental design, including a longer period of fasting, to generate enough hunger in these animals.

**Fig 4.**
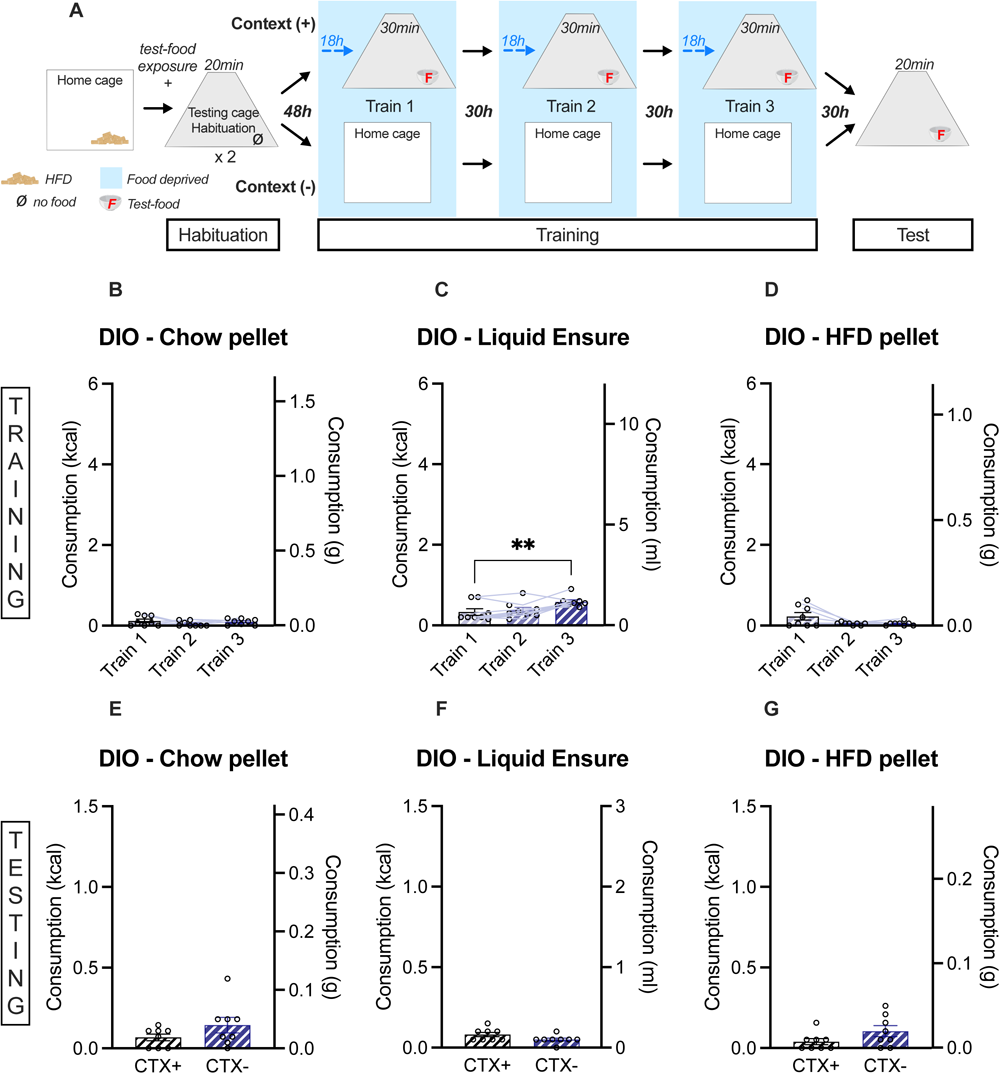
Diet-induced obesity (DIO) animals are not displaying context-induced overconsumption regardless of the reinforcers. (A) Experimental design of context-conditioned overconsumption paradigm. (B) Food consumption during training, while trained with chow or (C) liquid Ensure (50%) or (D) HFD (60%) expressed in kcal or grams. (E) Food consumption during testing while trained with chow or (C) liquid Ensure (50%) or (D) HFD (60%) expressed in kcal or grams. (B-D) n=8/group RM-ANOVA. (E-G) n=8/group t-test. **p<0.01.

## 4. Discussion

Overconsumption is a critical aspect of both clinical obesity as well as binge-eating disorder (Brown & James, 2023). Therefore, understanding the mechanisms that drive non-homeostatic feeding are critical. The conditioned overconsumption task specifically models one aspect of overconsumption, namely cue-driven eating. In our prior study we identified some parameters governing conditioned overconsumption, namely that spaced training was required, as well as multiple training sessions (Stern et al., 2020). We also found that a population of neurons in the insular cortex marked by Nos1 are required for the overconsumption effect (Stern et al., 2021).

Here we extended our findings by testing three main hypotheses. 1. That the task would be effective at eliciting overconsumption in both males and females. 2. That highly palatable food will also drive the effect 3. That experimentally obese mice will be prone to overconsumption in the task.

First, we find that the task works in both males and females. Exploring sex differences is important for pre-clinical work in which treatments may be tested (Shansky & Woolley, 2016). As binge-eating occurs in both males and females, it is important to have a task that is effective at inducing overconsumption in both sexes. Here we show that males and females learn the task equivalently and have an equivalent response.

Second, we find that sucrose is surprisingly not effective at inducing overconsumption compared to chow, Ensure and HFD. Similar findings were recently found in a fixed ratio operant training task, in which chow pellets were highly preferred over sucrose pellets (Karlsson et al., 2018; Karlsson & Cameron, 2023). However, during progressive ratio testing mice showed increased motivation to work for sucrose (Karlsson & Cameron, 2023). These results suggest that although mice prefer chow when given the option, they will work harder to obtain sucrose, although the underlying reasons for this discrepancy are not clear. In an appetitive Pavlovian task, sucrose also elicited more magazine entries than pellets (Rescorla, 1997). However, this testing was conducted in parallel and the order was not counterbalanced, thus this difference may reflect a reduction in neophobia over the testing period. Similarly to our findings with Ensure (a chocolate milkshake) and sucrose, strawberry milkshake was superior than a saccharin/glucose mixture in a touchscreen operant task (Phillips et al., 2017). We have not found any studies comparing HFD to other reinforcers, likely because it is challenging to put HFD into pellet form for operant training. However, because context-conditioned overconsumption is trained and tested with free access to solid food in cups or liquid substances in sippers, we can directly compare the caloric intake of all of the presented reinforcers.

In our studies we find not only a difference in consumption during training, but also during testing. We propose that there may be two explanations for this effect. First, chow pellets, Ensure and HFD all have a complete nutritional profile. Although Ensure is high in sugar, protein and fat, it contains the full profile of nutritional needs. Similarly, although HFD contains 60% fat, it still contains protein and carbohydrates to maintain a complete nutritional profile. In contrast, liquid sucrose contains only sucrose with no other additional nutrients, and sucrose pellets contain sucrose and dextrose, with no protein and minimal fat. Therefore, it is possible that complete nutrition is required to maintain learning of the task and drive the overconsumption response. Indeed, in preference tests, rodents will choose diets that contain protein over diets that do not, even if those diets are diluted or concentrated (Booth, 1974). The second potential explanation is that sucrose is less palatable compared to chow, Ensure and HFD. Compared to all three of these, mice did not eat significant amounts of sucrose during the training sessions, indicating that they do not have high baseline motivation for sucrose, even when fasted. Moreover, the addition of chocolate flavor to the sucrose pellets did not increase consumption during training or testing. However, sucrose is a very commonly used reinforcer for many operant-based training tasks (Reilly, 1999). It is therefore possible that sucrose is sufficient to maintain performance in food-restricted mice even if they don’t have a high baseline motivation to consume it, as was found in the case of progressive ratio testing (Karlsson & Cameron, 2023). In line with our hypothesis, we found contrasting results when we tested taste aversion learning: Ensure does not support CTA, whereas sucrose does. This supports the idea that Ensure is more appetitive than sucrose and thus the aversive association with gastric malaise is not sufficient to overcome the motivation to consume this substance.

One thing we noted was that even though Ensure robustly supported conditioned overconsumption learning, mice need to consume less Ensure during training compared to chow (when converted to kcal) to achieve that result. This is possibly due to the difference between liquid and solid food being offered, as other studies have shown that liquid reinforcers are superior (Kim et al., 2017). In addition, during the task, chow is offered without water available, but Ensure has significant water content, which can also satiate mice, leading to less caloric consumption.

When we tested HFD as a reinforcer, we first found no significant effect on overconsumption, which was largely driven by high amounts of consumption in the control group. This was perplexing, as it was previously demonstrated that HFD can support conditioned overconsumption (Mohammad et al., 2021). One key difference between the two studies is that Mohammad et al., used a sated control group, in which non-fasted mice are placed into the training context with food. Because these mice are not hungry, they tend not to eat in the training phase or the testing phase, as we showed previously (Stern et al., 2020). This led us to believe that perhaps mice were being habituated to HFD by increasing the number of exposures to it during training. In our case, we use CTX– mice as the control in order to control for the fasting period, but adding exposures to HFD in the homecage was able to reveal a similar effect of HFD on overconsumption.

We also found that when mice were not given additional exposures that HFD that we did not see a significant increase in cFos compared to mice given chow, although this effect was driven by differences in males and females (Supplementary Figure 4). The difference may be due to a difference in HFD sensitivity as it has been shown that females might be protected by estrogen (Acharya et al., 2023; Butler et al., 2020; Fabre et al., 2023).

Lastly, we tested whether DIO mice would be more prone to overconsumption in this task. DIO mice have been tested in Pavlovian and operant training paradigms, and have displayed deficits in motivation and learning, particularly in the progressive ratio task (Cordner & Tamashiro, 2015; Huwart et al., 2022; Kanoski et al., 2007; Sharma et al., 2013). In one notable exception, mice were trained to increase the level of force to obtain food rewards rather than number of operant responses (Matikainen-Ankney et al., 2023). In this case, DIO mice exerted more effort to obtain food rewards, suggesting that their lack of performance in the task is not due to low motivation, but rather due to a devaluation of delayed rewards. However, DIO mice have never before been tested in the conditioned overconsumption task to our knowledge. Interestingly, no matter which reinforcer we tested, DIO mice never consumed significant amounts during training, nor developed an overconsumption response. It is interesting, however, that these mice also do not eat HFD during training sessions, although they did escalate Ensure consumption over the three training sessions. This results might suggest that DIO mice require a longer fasting period to display the same hunger levels as lean mice, as other studies have indicated (Kanoski et al., 2007). In humans, one study showed that females with obesity are unable to form cue-reward associations with food, but they are able to form them with money (Zhang et al., 2014), although other studies have shown that individuals with obesity are more susceptible to cues that predict food (Polivy et al., 2011). These results might therefore also suggest that although food cues can induce overconsumption, different mechanisms are recruited to maintain weight in the case of high energy balance.

Overall, we find that context-conditioned overconsumption is more effective using complete nutrition of chow and highly palatable substances compared to sucrose, and that the task is not effective in DIO mice. Future studies are required to untangle whether these effects are due to postingestive effects, palatability or differences in motivation.

## Author Contributions

S.B. and D.L-S. contributed to conceptualization, investigation, formal analysis, visualization, writing – original draft, M.J.O.C, J.V. and Y-S.H contributed to investigation, S.A.S. contributed funding acquisition, project administration, supervision, conceptualization and writing-original draft.

### Abbreviations

CTX+: context +
CTX-: context
DIO: diet-induced obesity
h: hour
HFD: high-fat diet
mL: milliliter
kcal: kilocalories
g: grams

## Acknowledgments

The authors thank the Max Planck Florida Institute ARC, and in particular Colleen Neiner, for animal care. This work was supported by the National Institutes of Health Brain Initiative R00DA04749 (S.A.S), the Brain Research Foundation Seed Grant (S.A.S), the One Mind Foundation Rising Star Award (S.A.S.), the Max Planck Florida Institute for Neuroscience, and the Max Planck Society

## Ethical statement

All experiments were approved by the Institutional Animal Use and Care Committee at the Max Planck Florida Institute for Neuroscience (Protocol #20-007) in compliance with National Institutes of Health guidelines for the care and use of laboratory animals.

## Declaration of competing interest

The authors declare no competing financial interests.

## Data Availability

Data will be made available by the authors upon reasonable request.

## Appendix A. Supplementary data

The following is the Supplementary data to this article.

## Supplemental Figure Legends

**Supplementary Figure 1.**
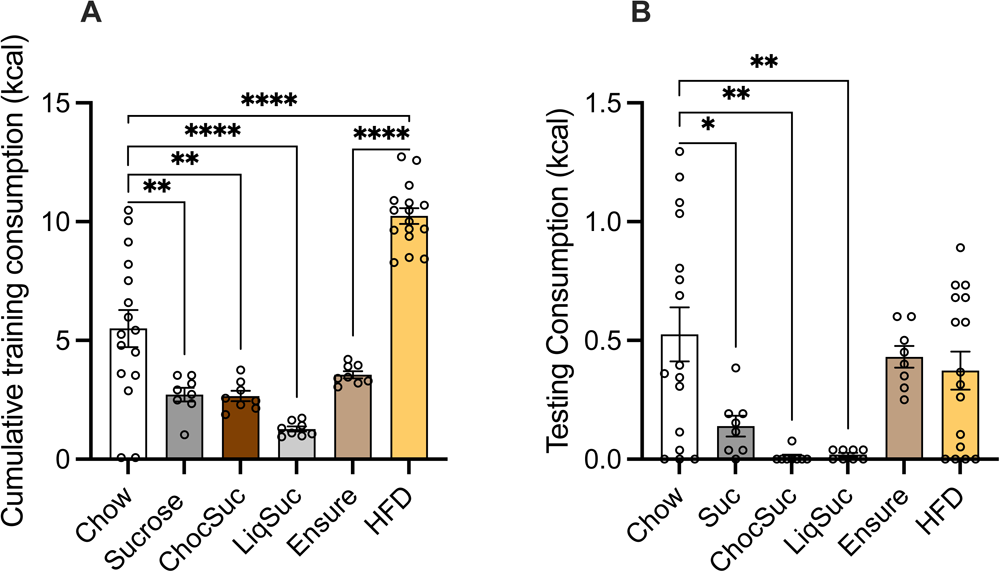
Inter-reinforcer analysis of training and testing food consumption. (A) Cumulative food consumption over the 3 training sessions for all type of tested food expressed in kcal. (B) Training food consumption for all different diets expressed in kcal. (A-B) n=16 for Chow and HFD, n=8 for Suc (sucrose), ChocSuc (chocolate sucrose), LiqSuc (10% liquid sucrose) and Ensure (50%), One-way ANOVA. *p<0.05, **p<0.01 and ****p<0.0001.

**Supplementary Figure 2.**
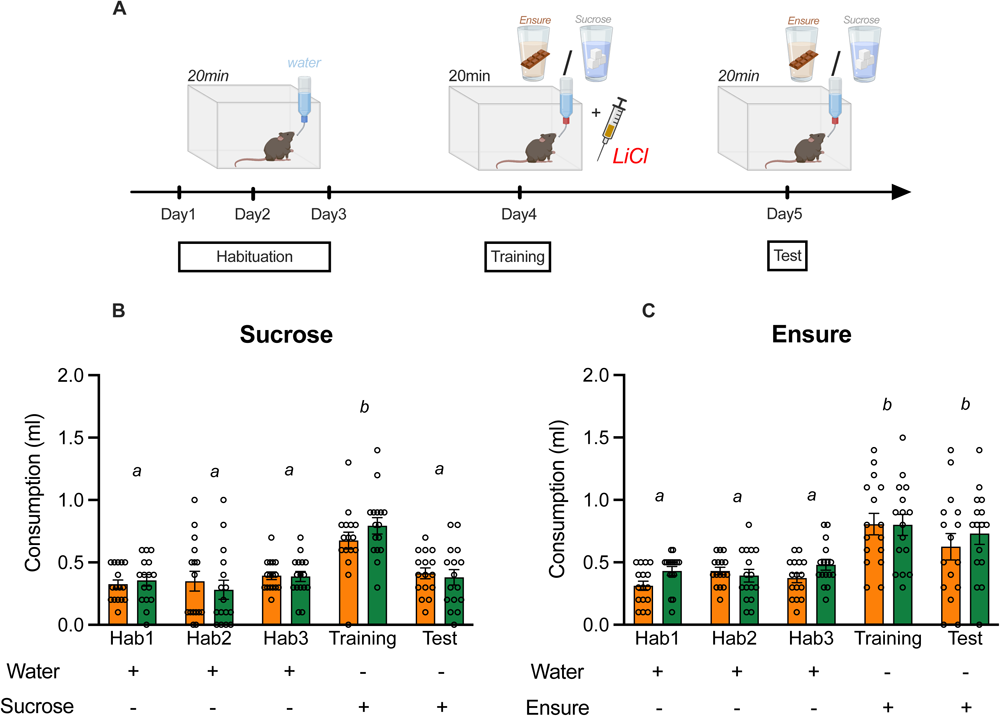
Detailed conditioned taste aversion (CTA) experiment. (A) Experimental design of CTA. (B) Water (Hab1-Hab3) or 10% liquid sucrose (Training and Test) consumption (mL) over the different sessions. (C) Water (Hab1-Hab3) or 50% Ensure (Training and Test) consumption (mL) over the different sessions. (B-C) n=16/group RM Two-way ANOVA. ‘a’s are statistically different than ‘b’s.

**Supplementary Figure 3.**
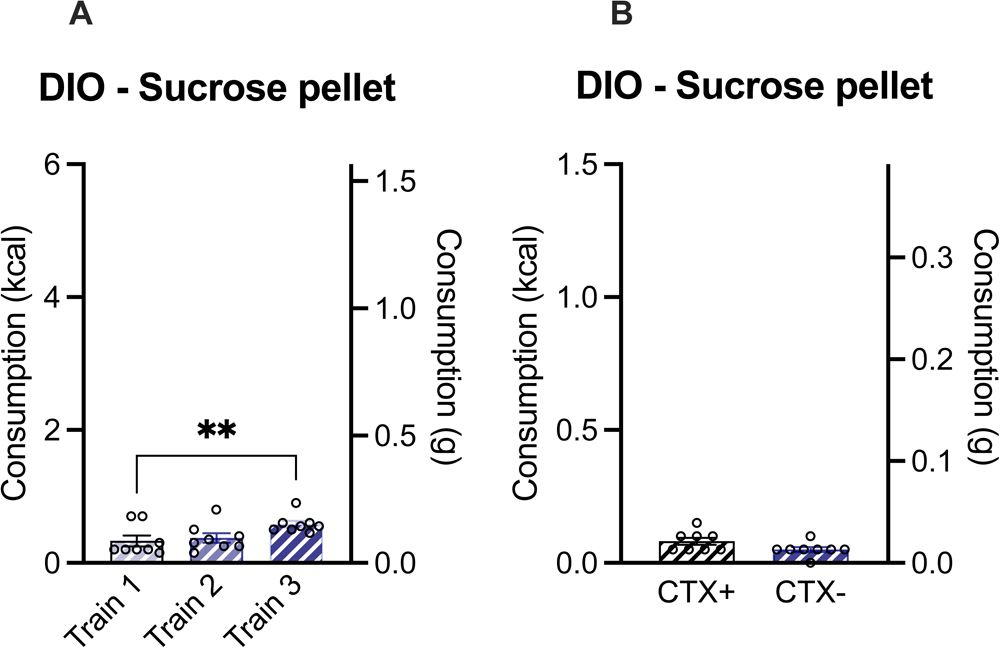
Conditioned-induced feeding using DIO animals and sucrose pellet as reinforcer. (A) Food consumption during training expressed in kcal or grams. (B) Food consumption during testing expressed in kcal or grams. (A) n=8/group RM-ANOVA. (B) N=8/group t-test. **p<0.01.

**Supplementary Figure 4.**
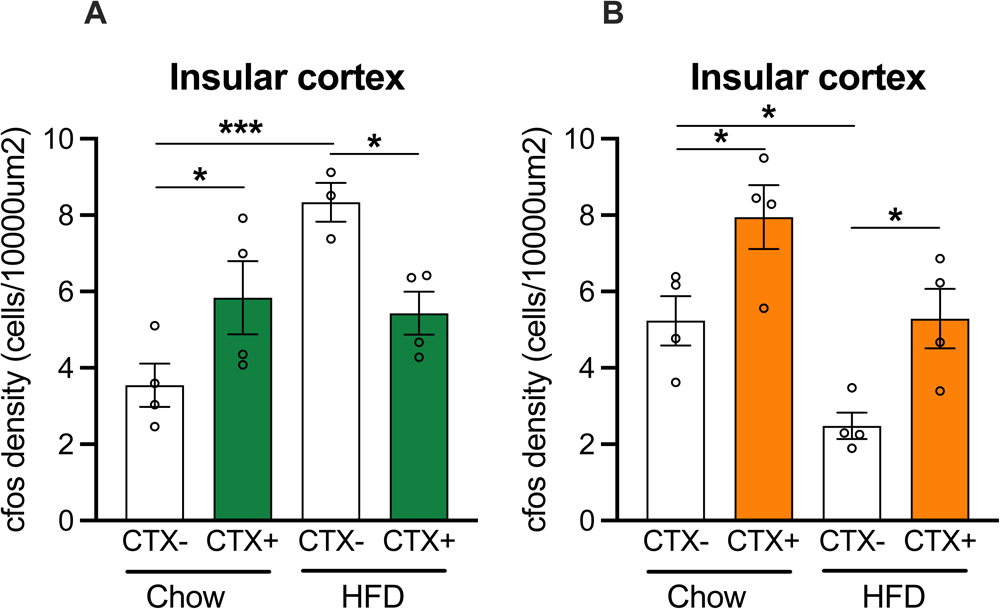
Post-overconsumption test cFos expression within females and males. (A) Quantification of cFos expression normalized by insular cortex area expressed in cells/10000μm2 for females and (B) males. (A) n=4 for Chow CTX+ and – and HFD CTX+. N=3 for HFD CTX-. Two-way ANOVA. (B) n=4/groups. Two-way ANOVA. *p<0.05 and ***p<0.001.

## Notes

### Competing Interest Statement

The authors have declared no competing interest.

